# Large-Scale Control of Neuronal Networks In Vitro Using Perforated Microfluidic Devices

**DOI:** 10.64898/2026.01.27.702069

**Authors:** Carl-Johan Hörberg, Jason P Beech, Ulrica Englund Johansson, David O’Carroll, Fredrik Johansson

## Abstract

Neurons in the brain are organized and connected into complex networks in which electrochemical signaling forms the basis for all brain function. Cortical neuronal networks are arranged in distinct modular, layered, and hierarchical structures, underlying its diverse functions such as learning, memory, or vision. Modern biotechnology has enabled an array of techniques to culture human neural cells, ranging from discreet co-cultures to complex developmental organoids, but all of which almost exclusively form unstructured and hypersynchronous networks. Overcoming this and capturing the functional and anatomical properties of the brain *in vitro* has proven to be a great challenge. Current techniques for guiding neuronal connectivity *in vitro* is often limited to a small fraction of the total population of neural cells, leaving most of the culture effectively unguided. To provide large-scale guidance of neurons in culture, we developed a microtunnel device which allows large-scale cell entry through an array of perforations, and guides neuronal network formation through a series of tunnels. Human neural stem cells capable of forming extensive neuronal projections were used to investigate several different microtunnel designs. One particularity noteworthy design which produced predominantly unidirectional growth was used to successfully validate its effect on propagation of neural activity on microelectrode arrays. Serendipitously, we found that our microtunnels had an extraordinary effect on signal-to-noise ratio and the quality of electrophysiological recordings with regards to number of active channels and detected spikes. Since we often found the neuronal growth surprising, we developed a simple computer model which could reproduce neuronal growth in the various tunnels, allowing computer aided design (CAD) of future projects.

## Introduction

Almost all *in vitro* neuronal cultures exhibit highly synchronous network activity with no obvious *in vivo* counterpart - a phenomenon which has been observed consistently for more than 40 years[15, 8]. Recent technological advances have greatly expanded the repertoire of in vitro neural models, particularly with regards to neurodevelopment and recapitulation of cellular phenotypes. Despite these advances, even the most elaborate in vitro cultures, whether as dissociated cultures[26], spheroids or organoids[40, 41], still display unrealistic synchronous network activity.

A large part of the function of the nervous system can be attributed to the connective topology of neurons[11, 29]. Simplified networks of neurons *in silico* have been shown to be able to exhibit complex information processing capabilities simply by virtue of their connective structure[38]. Perhaps in recognition of the importance of neural network architecture, mapping the ‘connectome’ of various animal neural networks has become a major frontier in neuroscience.

The recognition of the importance of neural connections strongly contrasts with the inaptitude of current technology to control neuronal connections in vitro. The homogeneous networks which are exhibited by most neuronal cultures, together with the absence of sensory input, are likely causes for the universal synchronized network bursts seen in cultures of neurons[27, 25, 4, 3, 20, 21]. While there’s been a lot of work done on addressing these issues, current techniques for controlling in vitro neuronal connections are still limited in several important ways. Generally speaking, different approaches trade off between throughput, directionality, fidelity, or electrophysiological access. The most common approaches are based on providing chemical or structural cues to direct growth[47, 2, 37, 48]. Techniques such as micro-contact printing can allow large-scale control of neurite outgrowth and cell positioning[28], but precise control of neuronal connections are mostly confined to a small number of neurons[46, 12]. Electrical fields can be used to control neuronal growth, but only with limited directional control[51, 52]. Other techniques such as 2-photon polymerized structures[1], electrospun fibers[13, 19], nanowires[18], microtopology[24] or other novel approaches have their own sets of challenges and have not yet demonstrated sufficient scalability, capacity for neuron guidance, or compatibility with electrophysiological techniques.

By far, the most common approach to controlling neuronal network formation in vitro has been to use various forms of microfabrication, particularly soft lithography, which is favored for its reproducibility and scalable production. Standard soft lithography is based on the casting of a polymer on a microfabricated relief substrate, creating an inverse copy of the microfrabricated mold[50]. This process will therefore yield a large bulk object with a fine imprint on one side. In microfluidics, the imprinted side is interfaced to a flat substrate, creating a narrow space of microtunnels with impermeable contact points between the polymer body and the substrate, which can be designed to almost any shape[49]. This approach can be used reliably to direct neuronal growth in many different ways[34, 35, 14], but is severely limited since neurons can’t be seeded into the narrow microfluidic compartment. Alternatively, cells can simply be seeded directly on the imprinted polymer surface, giving gross control of neuronal growth[31]. In this context, the fidelity of neuronal control and electrophysiological access is greatly reduced.

In this study, we aimed to overcome some of these challenges by creating a microfluidic system in which neuronal growth could be controlled with high fidelity on a large scale while also allowing electrophysiological access with microelectrode arrays. We used a 2-step approach to create perforated microtunnel devices that allow for large-scale control of neuronal networks on microelectrode arrays in vitro. Perforations allow for passive bulk entry of cells into the microtunnel compartment, which meant that cells could be seeded using protocols similar to those used to seed cells on traditional planar substrates. We tested several microtunnel designs aimed at promoting unidirectional growth. The observed growth patterns seldom matched our predictions, but it allowed us to develop a simple computer model to reproduce how neurons grow in these devices and thus allow for computer-aided design (CAD) of in vitro neuronal networks. One of our designs yielded a particularly unidirectional growth pattern and was applied to microelectrode arrays. Using human iPSC derived neurons, we could demonstrate that these microtunnels directed neural activity in one preferred direction. This simple design principle could be applied to create a multitude of different neural network topologies in vitro.

## Material and methods

### Microchip production and preparation

Perforated microtunnel devices were made by a two-step process by first creating a thin perforated PDMS membrane and subsequently attaching microtunnels. These two steps relied on casting PDMS on two different master molds, which were both created with photolithography on 4” silicon wafers in the following way. Wafers were dried at 200°C for 5 minutes and oxygen plasma treated (Plasma Preen, NJ, USA) for 60 s at 5 mBar.

For microtunnel structures, a ≈ 10µm layer of mrDWL5 resist (Micro Resist Technology GmbH, Berlin, Germany) was spun onto the wafer at 1100 rpm for 60 s. Pre-exposure baking was done by ramping up from 50 to 90°C over 20 minutes, and exposure was done on an MLA150 maskless lithography system (Heidelberg Instruments GmbH, Heidelberg, Germany). Post exposure baking was done the same as pre-exposure bake and development was done in Mr DEV 600 (Micro Resist Technology GmbH, Berlin, Germany) for 15 minutes plus 5 minutes in fresh developer followed by an IPA rinse and drying with nitrogen.

For perforated membranes, a 100 µm thick dry film resist (SUEX, K200, DJ Microlaminates, Sudbury, Massachusetts, USA) was laminated onto a 4” silicon wafer using a laminator (Catena 35, Acco UK Ltd, Buckinghamshire, UK) at 65°C. Pre-exposure baking was done at 85°C for 5 minutes on a hotplate (Model 1000-1 Precision Hot Plate, Electronic Micro Systems Ltd, West Midlands, UK). The wafer was exposed with 365 nm for 27 s at a lamp power of 30 mW/cm^2^ in a contact mask aligner (Karl Suss MJB4 soft UV, Munich, Germany) through a chrome mask (Delta Mask, Enschede, The Netherlands). Post-exposure baking was done at 85°C for 5 minutes and developed in Mr DEV 600 (Micro Resist Technology Gmbh, Berlin, Germany) for 15 minutes followed by an IPA rinse and drying with nitrogen.

In both cases, the resulting structures were hard baked at 180°C for 30 minutes. To reduce adhesion of PDMS, an ALD (Fiji, Plasma Enhanced ALD, Veeco, NY, USA) was used to deposit a layer of aluminium oxide (1 nm) followed by a monolayer of Perfluorodecyltrichlorosilane (FDTS).

The ‘pillar’ array was copied multiple times by casting PDMS, creating PDMS molds with 100 µm deep wells. These PDMS molds were heat-treated with 180°C for at least one our, after which PDMS could be cast and cured at, again, 180°C, without strong PDMS-PDMS bonding. This yielded arrays of PDMS pillars, which were used in subsequent stages to create perforated membranes. To create concave perforated membranes, these PDMS pillars were heat-treated at 180°C for at least one hour, after which a small amount of PDMS was spread by hand (wearing a nitrile glove), enough to fill just the space between the pillars, but not to cover them. These were then cured at 180°C for 5-10 minutes, after which a perforated membrane could easily be peeled off.

After the perforated membranes were created, microtunnels were added in a separate step. By applying a very small amount to the microtunnel master mold, and gently wiping off excess PDMS, PDMS accumulated only in the grooves of the master mold. The perforated membranes could then be applied and cured at 40°C over night. When peeling off the perforated membrane from the master, microtunnel shapes were now attached to the perforated membrane without clogging the perforations. In a few cases, a very thin film of PDMS accumulated on the patterned side of the perforated membrane, but these could easily be removed manually with fine tweezers.

To create ‘frames’, a 4-by-4 piece with perforations was cut out by hand before attaching the microtunnel pattern. This was placed in a hand-cut PDMS frame, and glued together with a small amount of PDMS, after which they had microtunnel patterns attached to them as described above.

To attach complete perforated membranes to microelectrode arrays, we aligned the perforations with the electrodes by hand, after lubricating the surface with pure ethanol. These assemblies were plasma-treated for 2 minutes, after which a small amount of poly-l-lysine (PLL) was added carefully, so as to only fill the microtunnels and not the upper surface of the perforated membrane. After one hour of incubation in 37°C, PLL was rinsed off and replaced with water, which was left to dry overnight. At least one hour before cell seeding, a small amount of laminin 10mg/ml dissolved in culture medium was added, again carefully filling only the microtunnels, and incubated in 37°C until cell seeding.

### Cell culture

A human neural stem cell line was provided to us by Prof. A. Björklund (Dept. Exp. Med. Sci., Lund University, Sweden). These cells were initially established by L. Wahlberg, ^°^A. Seiger, and colleagues at the Karolinska University Hospital, Stockholm, Sweden.

Neural stem cells were expanded as neurospheres in DMEM/F12 (Invitrogen, UK), with 2.0 mM L-glutamine (Sigma-Aldrich, USA), 0.6% glucose (Sigma-Aldrich, USA), N2-supplement (Invitrogen, UK), 2.0 *µ* g/mL heparin (Sigma-Aldrich, USA), 20 ng/ml human basic fibroblast growth factor (Invitrogen, UK), 20 ng/ml human epidermal growth factor (PROSPEC, Israel), and 10 ng/mL human leukemia inhibitory factor (PROSPEC, Israel).

Neural stem cells were passaged roughly every 14 days by dissociation with accutase (Thermo Fisher Scientific, USA) where cell count was assessed by trypan blue exclusion. After dissociation, cells were either seeded for an experiment, or reseeded for further expansion. Passages between 14 and 16 were used in our experiments. Neural stem cells were seeded and cultured in medium containing (DMEM/F-12 medium with 2.0 mM L-glutamine, 0.6% glucose, N2-supplement, 2.0 *µ*g/mL heparin and 1% fetal bovine serum (Invitrogen, UK)) by pipetting a 10*µ*l droplet containing 100’000 live cells.

Commercial human iPSC-derived neurons and astrocytes (FUJIFILM Cellular Dynamics, USA) were seeded in a 1:6 ratio, according to suppliers instructions, to a total of 100’000 cells per MEA. These cells were grown in BrainPhys (StemCell Technologies, Canada) neuronal medium, with N2, Penicillin, Streptomycin and the manufacturers associated cell supplement, neural supplement B and nervous system supplement (FUJIFILM Cellular Dynamics, USA).

### Immunocytochemistry

The samples were fixed for 10 minutes with 4% paraformaldehyde solution, after which the samples were rinsed with PBS. Samples were then pre-incubated in blocking solution (PBS, 1% BSA and 0.1% Triton X-100) for 30 minutes. All antibodies were diluted in blocking solution. *β*-III-tubulin-reactive mouse antibodies (Thermo Fisher, USA) were used at a 1:2000 the original concentration to stain neurons. The samples were incubated with primary antibodies for at least two days. Mouse-IgG-reactive donkey antibodies, with a conjugate AF488 fluorophore (Thermo Fisher, USA) was diluted 1:200. The secondary antibodies were incubated for at least two days.

### Image acquisition

Images were captured with a Zeiss Observer Z1 (Zeiss, Germany) inverted epifluorescence microscope with a motorized stage. All samples were entirely photographed as large tiled images with a Hamamatsu Orca-Flash4.0 (Hamamatsu, Japan). Images were acquired and stitched using Zeiss Zen 3.9.

### Image analysis

Tracing neurites was achieved by manually tracing identified neurite terminus, and back-tracking neurites from identified neurite tips. This way, complete neuronal morphologies were rarely attained, but rather different size sections of neurites. In some cases, it was easy to identify that a neurite was originating from a certain cell soma. Statistical estimates of confidence intervals could be achieved by bootstrapping means, by resampling with replacement the entire population of neurites 6000 times.

### Electrophysiological recording

Electrophysiological recording was done using a MEA-1060 (Multichannel Systems, Germany) recording system, at least once per week. Recordings of spontaneous activity were done at 32000Hz for 10 minutes after an initial 5 minutes of equilibrating at room atmosphere.

### Spike detection

All data analysis was done in python and C. Raw signal was filtered with scipy using a Butterworth band-pass filter with low-cut of 250 Hz, and high-cut of 10 kHz. We detected spikes by using a custom C-script which included any event which exceeded +6 or -6 standard deviations of the estimated noise.

### Signal-to-noise ratio and active channels

All channels which exhibited more than 12 spikes per minute was defined as active. In order to calculate noise, we looked at all active channels, and estimated noise as periods in the recording that was at least 2 ms away from a spike. To estimate signal, we took the peak absolute value of each spike, stored this value for all spikes in a vector, and measured the signal as the threshold for the top 1 percentile of spike amplitudes. This was repeated for each channel.

### Transfer entropy

Transfer entropy was calculated by first postulating that at any given point in time, a channel can either exhibit a spike or not. From this, we constructed joint probability matrices of *pX*_*t*=*i*_; *pX*_*t*=*i*−*l*_ and *pX*_*t*=*i*_; *pX*_*t*=*i*−*l*_; *pY*_*t*=*i*−*l*_. Where *pX* and *pY* is the probability that a channel *X* or *Y* respectively, exhibited a spike. From this, the transfer entropy is given by *T*_*y*→*x*_ = *H*(*X*_*t*=*i*_|*X*_*t*=*i*−*l*_) −*H*(*X*_*t*=*i*_|*X*_*t*=*i*−*l*_, *Y*_*t*=*i*−*l*_). This was done for a range of *l*, up to a corresponding *l* of 10 ms. This was done for all recordings of PDMS cultures after 37 days of culture.

### Simulation of neurite growth

The simulation was written in Python using numpy. Microscopy images of microtunnel designs were used to manually create binary bitmaps representing pathable and non-pathable coordinates. A neurite tip was considered as similar to a rigid body with momentum, and its position was updated with a fixed time step, but moved by a fixed distance for each step.

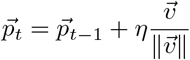

*p* is the position of the neurite tip, which at time *t* is updated by moving a fixed distance *η* in the normalized direction of its velocity 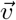. The velocity was updated based on a few factors described below.

The neurite tip velocity was continuously increasing radially outward from the starting point. The neurite tip ‘senses’ edges which lie within a radius around it. By using a simple flood-filling algorithm, the algorithm measures the shortest path to all edge coordinates, such that the neurite tip can’t sense edges which are behind obstructions.

The neurite tip is both attracted by distant edges, while being repelled by close ones. If the neurite tip attempted to move to an unpathable coordinate, the velocity was deflected by using the approximate normal of the edge.

The response to edges was only considered if they were with a given cone in front of the neurite tip. The ‘front’ of the neurite tip was not the same as the direction of the velocity, but an average of the velocity over a given number of time steps, which we called the neurite polarity.

The neurite itself was always considered as a straight line between the origin of the neurite and the tip. However, when this straight line inevitably passes over an unpathable coordinate, this contact point becomes a new origin for the neurite, until the neurite tip navigates to a point which again lies within an unobstructed line to the previous origin. This new origin also served as a pivot point where the angle between it and the previous origin was used to calculate ‘neurite stiffness’ by adding velocity to reduce this angle.

Additionally, the neurite tip velocity was given a small random velocity, and the velocity was also dampened by a percentage of its total velocity, each iteration. All parameters mentioned above (search radius for edge detection, angle for edge detection, and their respective attractive force, etc.) were tuned manually until the simulated growth matched that of the measured.

### Experimental design

Human neural stem cells were used for evaluating growth patterns, while human iPSC-derived neurons were used for testing electrical activity. Human neuronal stem cells were seeded with various microtunnels on three separate occasions. Tracing of neuronal morphologies was done on all identifiable neurons from at least three cultures per microtunnel design from across these replicates. In total, we traced 74, 119, 74 and 56 neurites in the four respective microtunnels as seen in figure 3E-H. In total, we seeded co-cultures of human iPSC-derived neurons and astrocytes (15% astrocytes, and a total of 100’000 cells per culture) on 8 “frameless” microtunnel cultures on MEAs, and 10 “framed” microtunnel cultures. Human iPSC-derived neurons and astrocytes were also seeded in MEAs without microtunnel devices, with 7 cultures of 15% astrocytes and 6 with high (50%) astrocytes.

## Results

### Microfluidic design

Our goal with this study was to control neuronal circuit formation across an entire population of neurons. We realized that a perforated design (Figure 1A) could be capable of producing such controlled networks. For guidance of neurites, we created several different microtunnel designs, but which all relied on a similar principle: obstructing or permitting neurite growth by means of physical constraint (Fig 1B). An array of perforations (Fig 1C) allows cells to enter the microtunnels (Fig 1D) in great numbers, which is otherwise severely limited using similar techniques. To further concentrate cell deposition, small sections of perforations could be added to a larger frame (Fig 1E), creating a larger basin where cell suspensions could be applied (Fig 1F).

**Figure 1.**
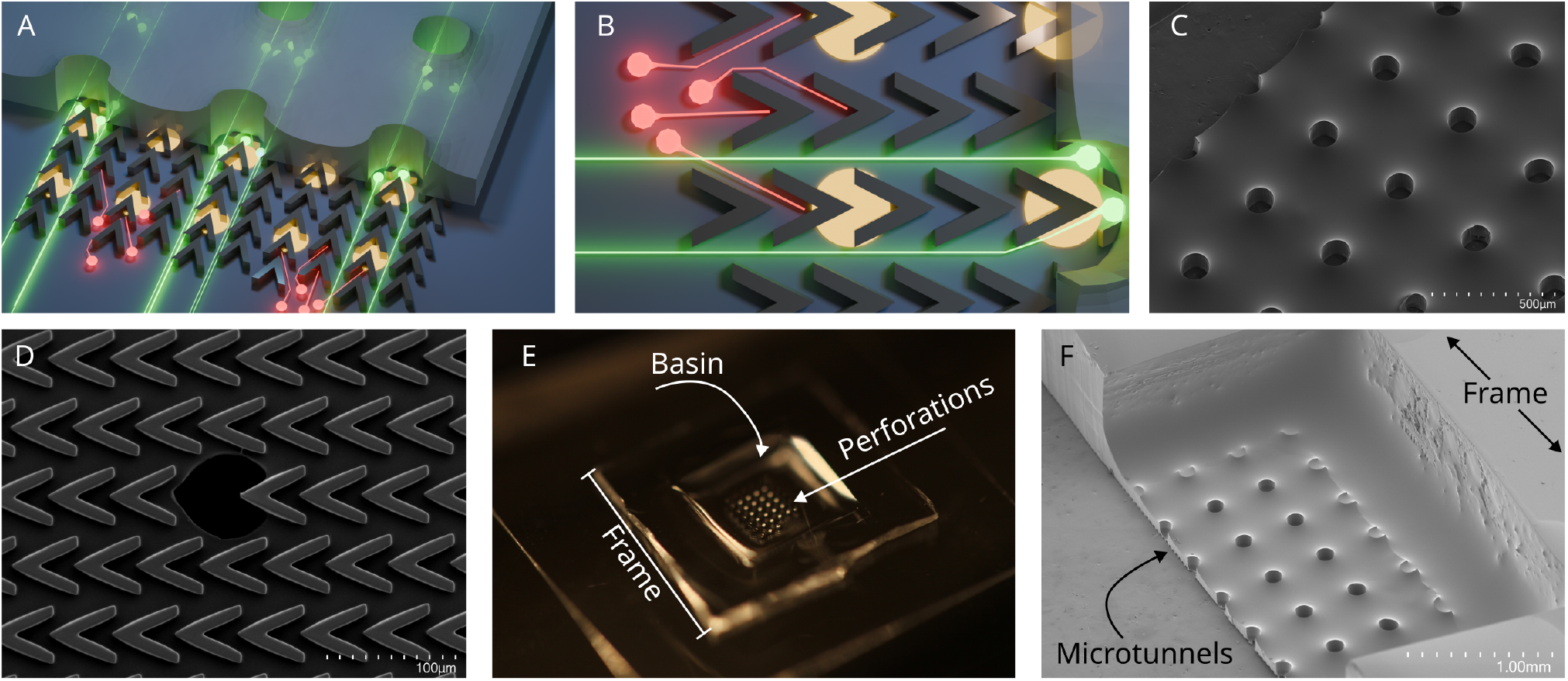
Experiment concept and device design. A) 3D Sketch of the layered structure of the device. The complete design consist of an upper relatively thick (100 µm; image not to scale) membrane with convex perforations. Under this layer lies a much thinner (10-20 m) layer of (in this case) V-shaped microtunnels. Neurons can be seeded from above, fall into the perforations, and grow into the microtunnels which lies between the perforated layer and the substrate below. The substrate below can be a microelectrode array; electrodes are shown here as glowing yellow discs. B) 3D sketch. Same as A, but seen from above. Here we also show the expected effect the microtunnels has on neurite growth, with red neurons showing how “backward” growth is inhibited, while green neurons showing how “forward” growth is permitted. Perforations are matched to electrode spacing so that each electrode is situated directly underneath a perforation. C) Electron micrograph of membrane perforations seen from above. D) Electron micrograph of one single membrane perforation seen from below, showing the tunnel structure underneath. E) Photograph of complete device. Here, a smaller perforated structure is welded to a larger and thicker ‘frame’ creating a small indentation to concentrate cells on the active electrode area of the microelectrode array. F) Electron micrograph of a fully assembled device cut vertically to show a cross-section of the full device. The frame is significantly thicker than the thin membrane which lies in the bottom of the indentation

A simple straightforward way of making perforated microtunnel devices was by employing a two-step process (Fig 2A) where perforations (Fig 2 C) were initially made by spreading a small volume of PDMS onto pillars (Fig 2D), and onto which micro-tunnels (Fig 2E) were added in a separate step. This circumvents the problem which otherwise faces this technique - that cells can’t be seeded directly into the microtunnel compartment (Fig 2F). The perforations allow for large-scale passive entry of cells into the microtunnel compartment, simply by seeding them onto the perforated membranes (Fig 2G).

**Figure 2.**
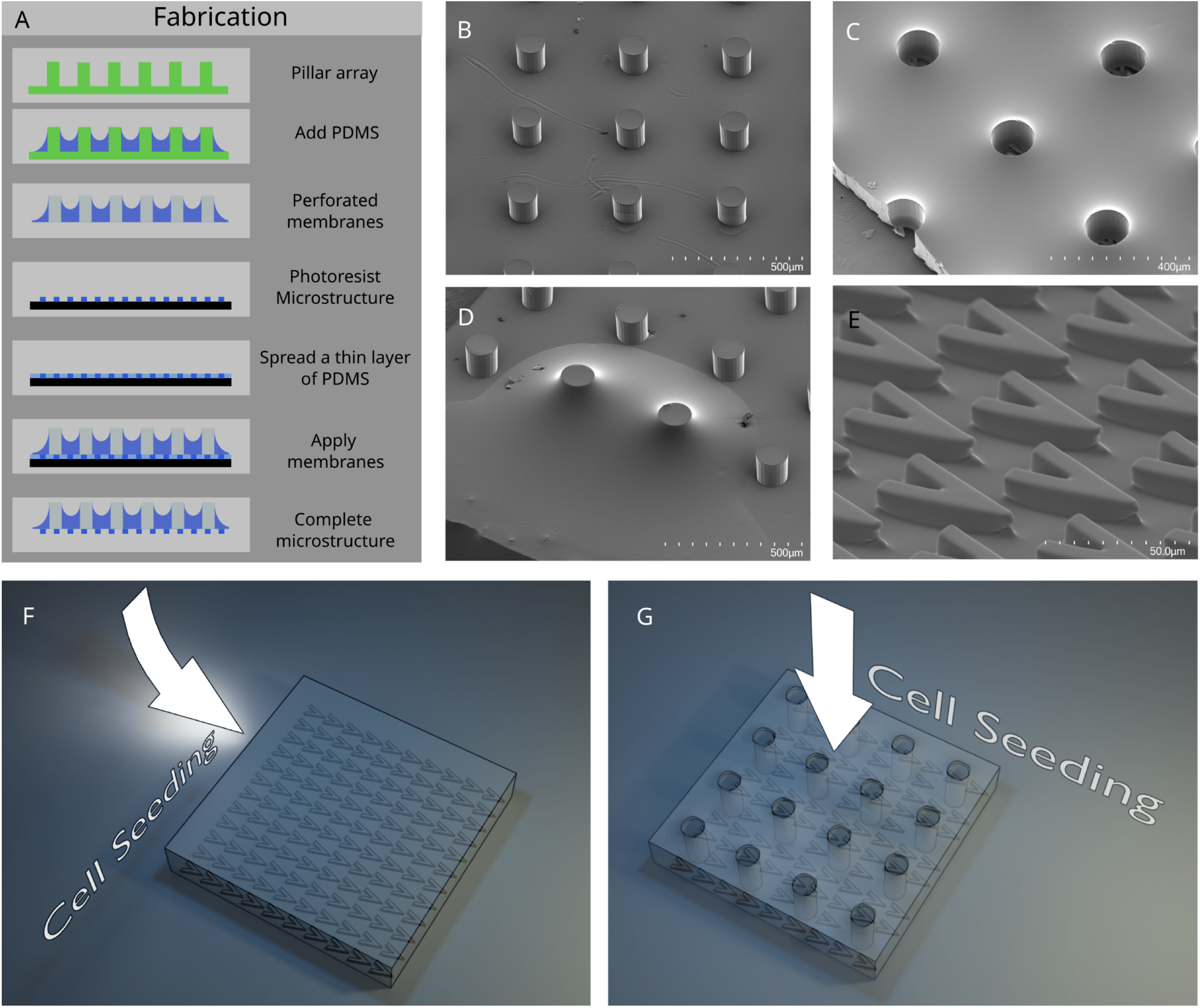
Device manufacture process. A) An array of pillars are first used to make a perforated membrane by simply applying an amount of PDMS to the pillar array that does not cover the pillars with PDMS. Then, by spreading a small amount of PDMS on a photolithographic mold, it’s possible to attach any microtunnel design to the perforated membranes. B) Scanning electron micrograph of a pillar array (PDMS copy). C) Scanning electron micrograph of a complete device seen from “above”, small parts of the underlying microtunnel pattern can be seen in the bottom of the holes. D) Scanning electron micrograph of PDMS being applied to pillars. E) Scanning electron micrograph of an example microtunnel design. F) Sketch of an ordinary microtunnel device, where only a small number of cells will enter the microtunnel compartment from the edges of the device. G) Sketch of complete perforated device, where cell seeding is possible by applying cells from above. Cell entry into the microtunnel compartment is facilitated by the perforations.

A potential concern with this design was that some cells could become trapped on top of the membrane, decreasing yield of cells inside microtunnels but also potentially adhering and forming an unguided network which could interfere with the purpose of the device to guide neuronal network formation. We deliberately made perforated membranes with a ‘convex’ shape (Fig 1 C), which we reasoned could discourage cells from growing into the perforations if they would happen to reside on top of the membrane. Using similar techniques, it was also possible to create ‘dimpled’ and ‘flat’ membranes (See Appendix 1).

The perforated membrane was designed for compatibility with microelectrode arrays (Fig3A), with perforations aligned to the electrode grid (Fig3B). For assessing different microtunnel patterns and chip designs, we utilized a human neural stem cell line, which is highly proliferative and migratory and forms glial- and neuronal cells in differentiation conditions[42]. When seeded, cells simply accumulate in the perforations by force of gravity (Fig3C), and migrate or grow (depending the migratory propensities of the cell type) out into the microtunnels. The architecture of the neuronal network which subsequently forms is now, apart from within the small perforations, dependent on the microtunnels and the growth patterns it permits (Fig3D).

**Figure 3.**
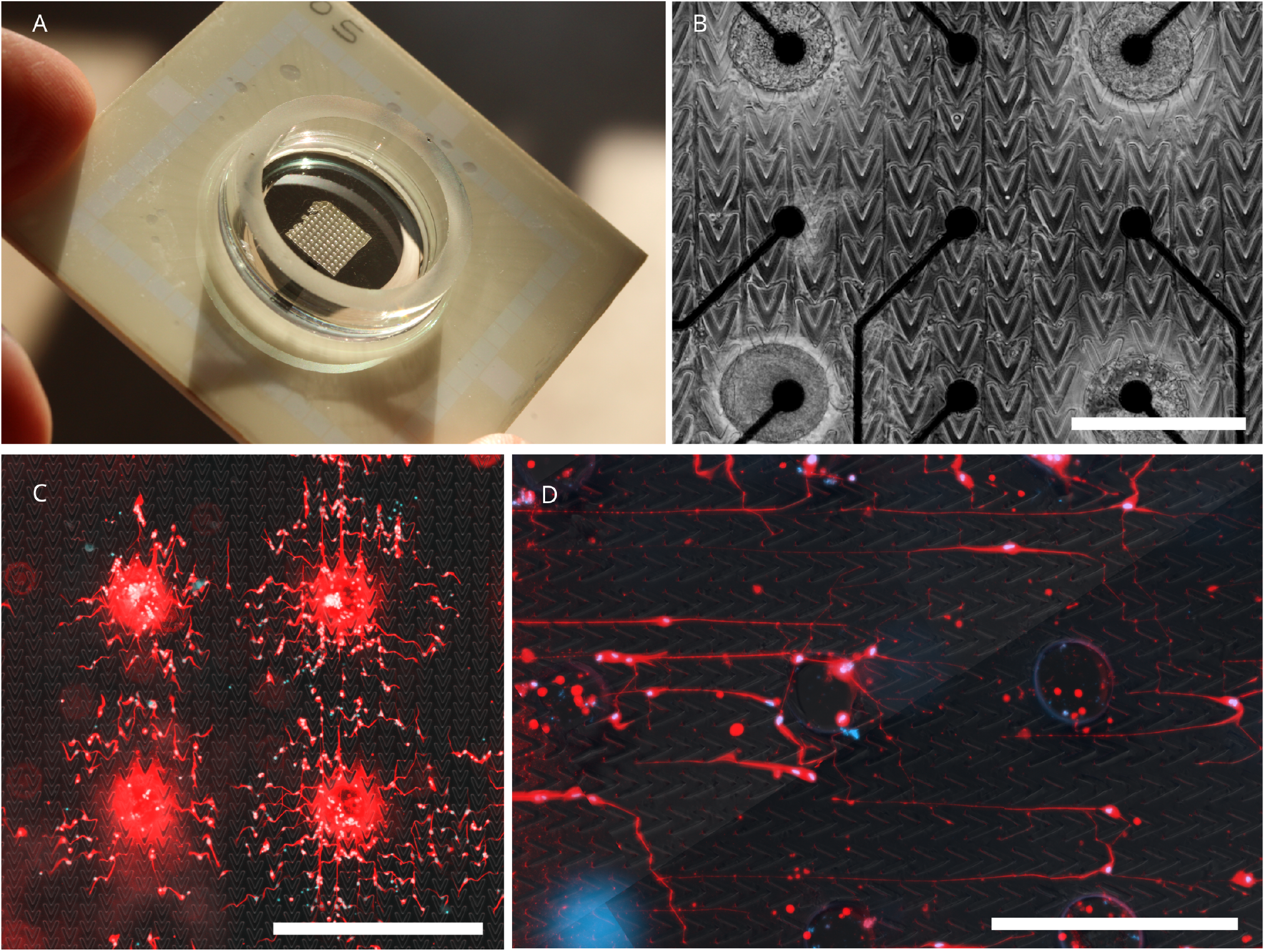
Device utilization A) Photograph of a frameless perforated device assembled on a micro electrode array. B) Phase contrast micrograph of iPSC-derived neurons growing on a device assembled on a micro electrode array. An abundance of neural projections can be seen extending along the vertical axis between the perforations. C) Epifluorescence micrograph of fixed and fluorescently labeled human neural stem cells 4 days after seeding. We see an initial rapid migration with cells migrating from the perforations and into the microtunnels. D) Epifluorescence micrograph of fixed and fluorescently labeled human neural stem cells two months after seeding. Cell at this age develop a more neuronal morphology, showing long (hundreds of micrometers) projections, showing clearly what effect the device has on the orientation of the projections.

### Different microtunnel designs produces unique growth patterns

We created 8 different microtunnel designs in total, all of which were intended to produce unidirectional growth. We took inspiration from established procedures but also tested several novel shapes. By seeding human neural stem cells on non-perforated versions of these microtunnels, we could evaluate their effects on neuronal growth by using fluorescence microscopy. When seeded, these cells do not form long neuronal projections (Fig 3C) but after 2-3 month in culture, most cells extend long projections and express *β*-III-tubulin. These cells were then fixed and stained using immunocytochemistry, and it was possible for us to manually trace individual neurites. Out of the 8 microtunnel designs, 4 were deemed interesting for further study and were analyzed thoroughly. All microtunnel designs that were tested produced highly distinct neurite growth patterns (Fig 3A-H). Several microtunnels produced dominant “forward” growth, but often produced an unwanted and unexpected “backward” growth as well. One microtunnel stood out as highly selective for unidirectional growth, sometimes growing almost 1 mm in one direction (Fig3A). This microtunnel design was therefore selected for further studies.

### Growth patterns can be replicated using a simple computer simulation

Since all microtunnel designs were intended to produce unidirectional growth, but often found paths that we did not expect when designing the microtunnels, we wanted to see if it would be possible to simulate the growth of neurons and thus simplify future attempts at designing neuronal networks in vitro. We wrote a simple python program which simulated the neurite growth tip as an inert body, with several different forces acting on its momentum. Using microscopy images of the different microtunnels, we created bitmaps which represented pathable and non-pathable coordinates. Neurite tips were given a random starting location and growth direction, and iteratively moved based on its current momentum (see methods for a complete description of the model). Briefly, the neurite tip was pushed away from its starting point and away from nearby unpathable regions, while being attracted by distant edges between pathable and unpathable regions.

The neurite was considered as a straight line between the starting position and the mobile neurite tip until an obstruction appeared anywhere along this straight line. This point of obstruction was then used as a pivot point (Fig A), the angle at which was used to implement ‘neurite stiffness’, adding resistance to bending this angle too sharply. This last step was crucial in recreating the criss-cross backward growth shown in Figure B and C.

A few hundred neurites were simulated per micro-tunnel pattern and were compared with the experimental data. Initially, we saw that most of the dominant growth patterns could be reproduced (FigB-C), but in order to match the relative distributions of these growth patterns, we manually had to tweak the different parameters until all different patterns matched their experimental counterparts. With few exceptions, the simulated data came very close to the experimental data.

### Microtunnels control propagation of action potentials

One microtunnel design stood out as preferably guiding neurites in one direction (Fig4A,E). Since we expected this microtunnel design to therefore support the most identifiable effect on the propagation of action potentials, we used this design for further testing. We manufactured perforated devices with this microtunnel pattern (Fig1D) and attached these to microelectrode arrays (MEAs). Commercial human iPSC-derived neurons and astrocytes were co-cultured on normal MEAs, and on MEAs with perforated microtunnels. We tested both microtunnel designs with (PDMS+frame) and without (PDMS-frame) “frames” (Fig 1E,F). The cultures were recorded once or twice a week for at least a month(Fig7 A), which allowed us to compare the development of spontaneous neuronal activity to standard culture conditions. Notably, the number of active channels and spike-rates of cultures with PDMS -in particular in PDMS+frame cultures-was much higher than in reference cultures (Fig 7 B-C).

**Figure 4.**
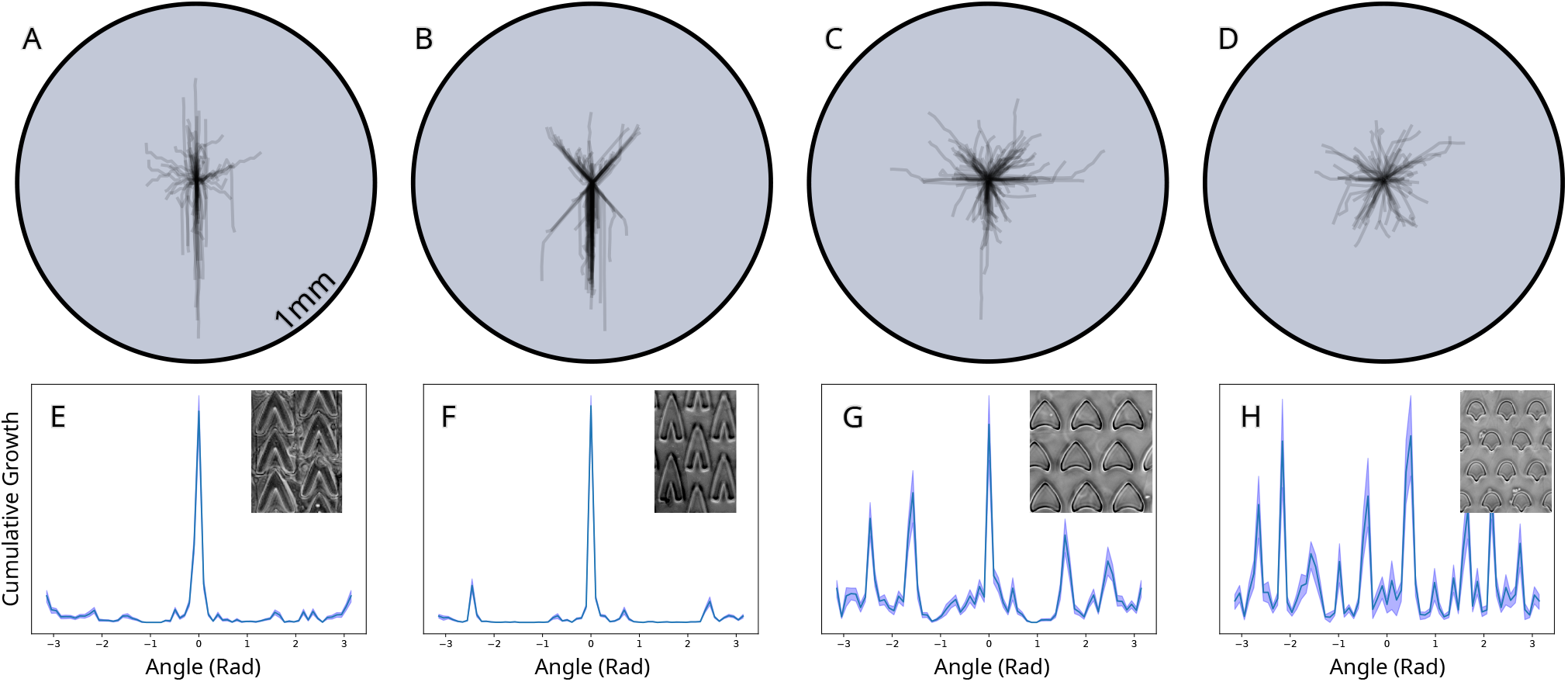
Reconstructed neurites from different microtunnel designs. A-D) All traced neurites plotted on a common origin. Colors indicates density of overlapping neurites. E-H) Normalized cumulative neurite growth for different microtunnel designs.

**Figure 5.**
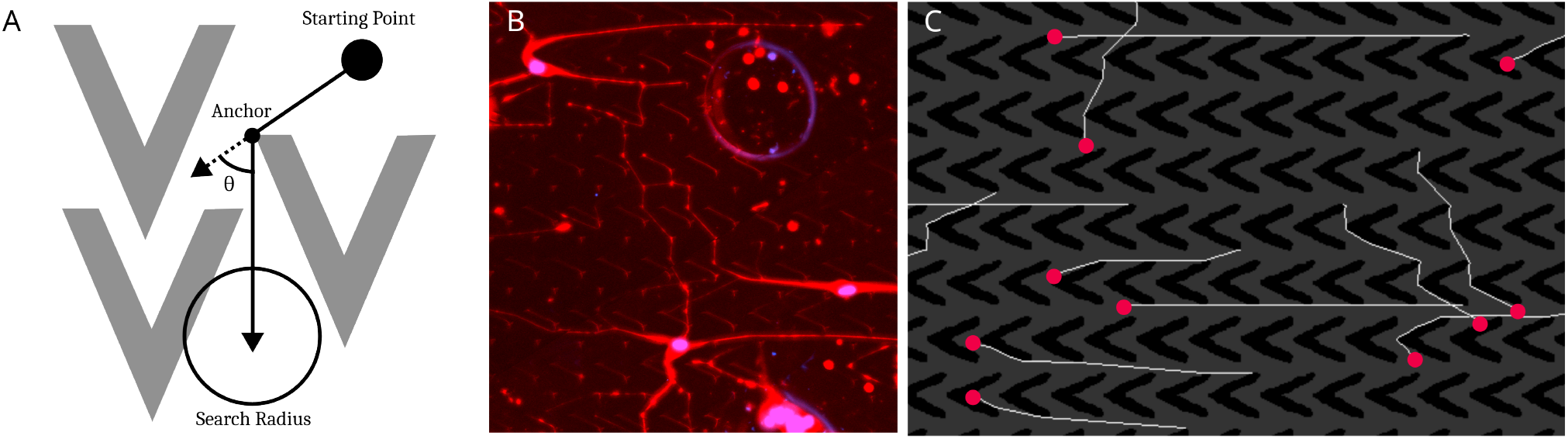
Simulation of neurite growth in different microtunnel designs. A) All parameters used in the simulation. A microscopy image from a microtunnel was used to create a binary bitmap indicating where a neurite is allowed to grow. The neurite tip moves as a rigid body with some random acceleration. The neurite has a polarity which is a weighted average of its past momentum, and searches for pathable area and edges of pathable area, and is attracted by both. The neurite is constantly pushed away form its last fixed point. Neurite stiffness is also implemented. B) Examples of simulated neurite paths. C) Examples of experimental neurite paths.

**Figure 6.**
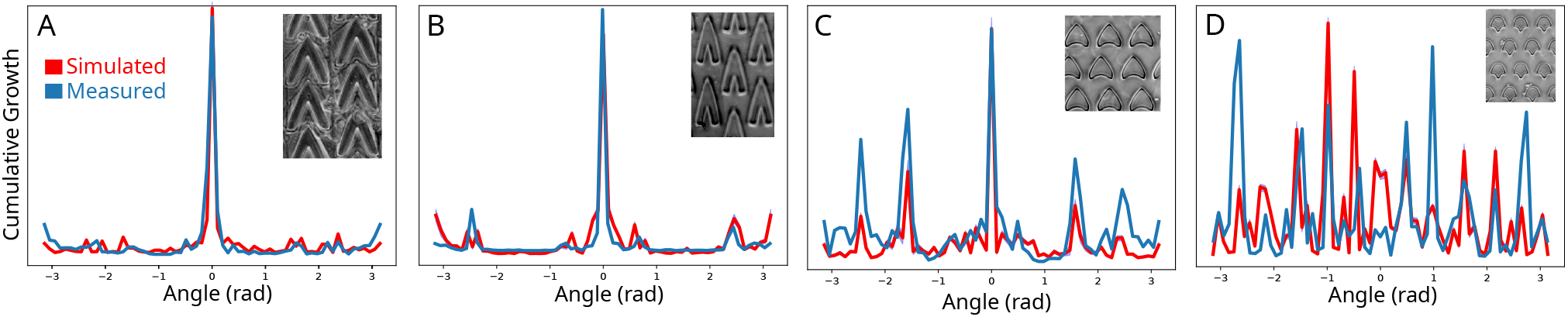
A-D) Comparison of experimental and simulated neurite growth probabilities for the different microtunnel designs.

**Figure 7.**
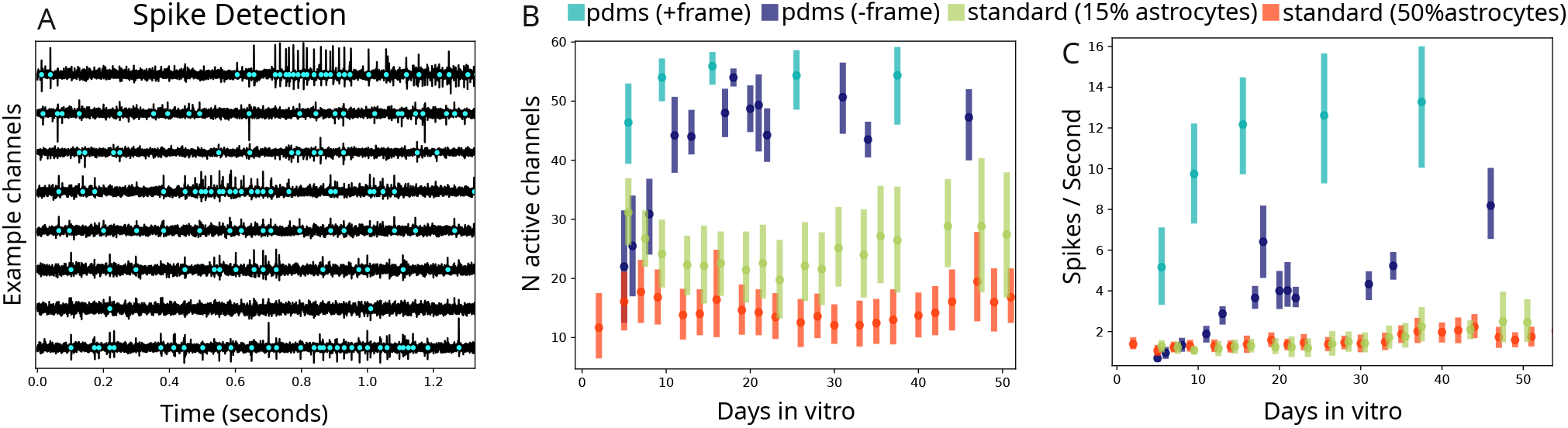
Electrophysiological analysis of neuronal development. A) Bandpass filtered electrophysiological data from 8 example channels, out of a total of 59 channels per MEA. This recording was taken from a PDMS+frame culture at 25 days in vitro. Teal dots indicates detected spikes. B) Number of active channels for PDMS+frame (teal), PDMS-frame (blue), standard culture (no PDMS guide) with 15% astrocytes (green) and a standard culture with high (50%) astrocytes (orange). C) Same as B but with spike-rates for each culture. Bars indicate 95% confidence intervals, while dots indicate means.

Upon closer inspection of electrophysiological data, we found an abundance of spikes occurring in very close temporal succession on adjacent electrodes (Fig 7 A). In addition, the spike amplitudes were much higher, at least three times the normal signal-to-noise ratio observed in standard cultures (Fig 8 B). The excellent signal, the high number of active channels and the large number of spikes provided the opportunity to do quantitative measurements of network connectivity.

**Figure 8.**
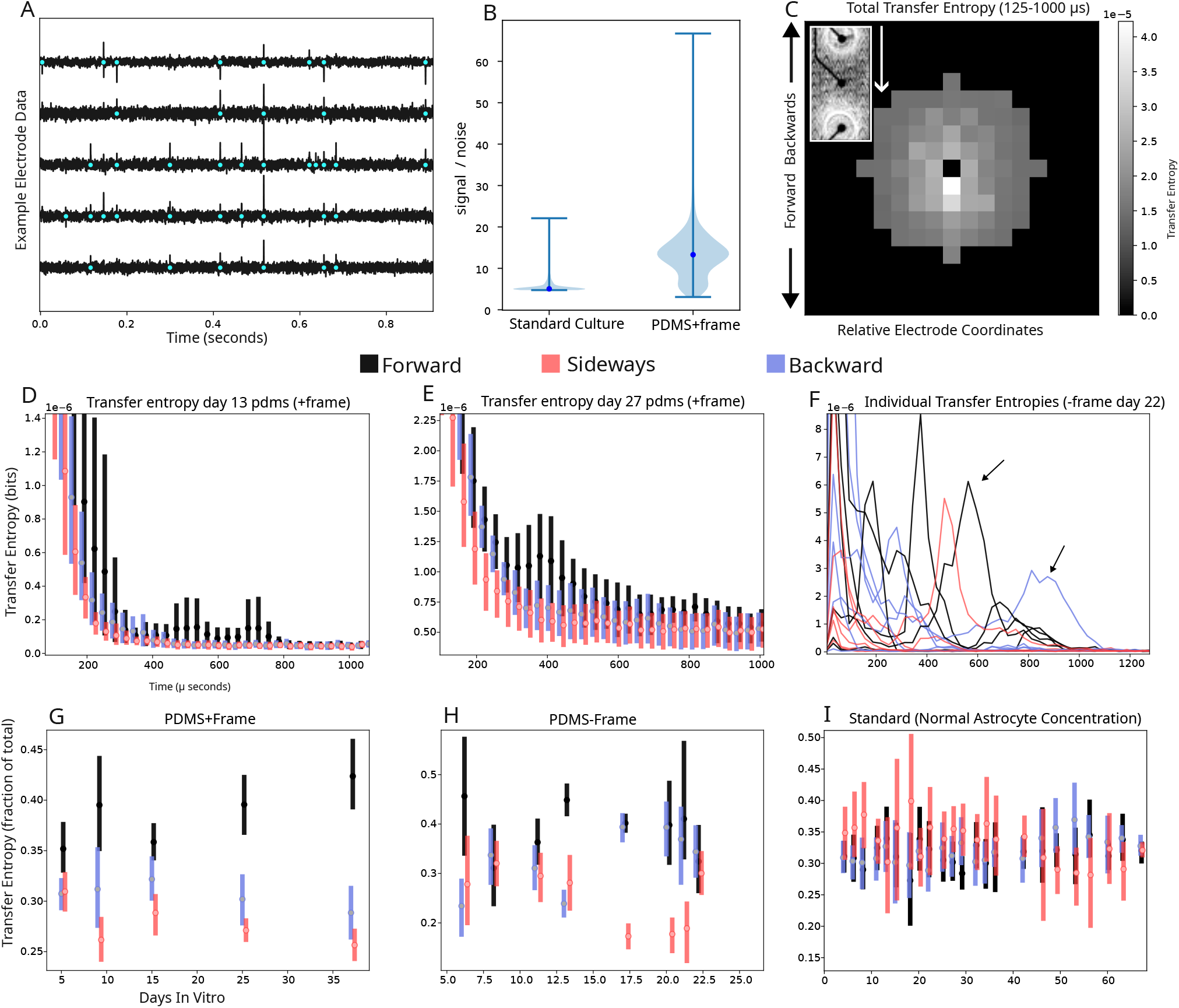
Measurement of neural activity propagation using transfer entropy. A) Bandpass filtered data of 5 example electrode channels, showing several spikes which occur in very close succession across multiple channels. Signal-to-noise ratio of PDMS cultures and standard cultures. C) Total transfer entropy of all PDMS+frame cultures at 37 days in vitro. Transfer entropy is summed over 125 to 1000 µs, and displayed as relative spatial electrode coordinates, such that pixels below the black pixel in the middle represents electrodes which are displaced in the direction which exhibited dominant growth by this pattern, which we call the ‘forward’ direction. This indicates that spikes exhibited by any electrode will in general increase the likelihood of spikes occurring on forward electrodes. D) Means and 95% confidence intervals of all PDMS+frame cultures and their respective forward (black), backward (blue) and sideways (peach) transfer entropy displayed as a function of time. E) Same as C but for PDMS+frame cultures at 27 days in vitro. F) Transfer entropies of all PDMS-frame cultures at 22 days in vitro, showing that individual cultures display disparate peaks of transfer entropy in different directions and at different times. G) The ratio between forward, backward and sideways transfer entropy and the total transfer entropy, showing confidence intervals and means which suggest a strong preference for ‘forward’ propagation. H) Same as G but for PDMS-frame. I) Same as G and H but for standard cultures with normal astrocyte concentration.

To test whether the network activity was influenced by the microtunnels, transfer entropy was measured from recorded spike trains. Transfer entropy is a non-parametric information theoretic approach which is often used to analyze neuronal connections[33]. By measuring transfer entropy between all electrode pairs, we could discern a clear directional preference (Fig8C). This suggested that neuronal activity indeed propagated unidirectionally, as one would suspect given the unidirectional growth induced by this microtunnel pattern (FigD).

Transfer entropy was consistently higher in the ‘forward’ direction for several hundreds of microseconds (Fig 8 D-E), but when looking at individual transfer entropies, it’s clear that all individual cultures (and possible electrode pairs) have strong individual peaks which do not coincide in time. Instead of looking at total transfer entropy in various directions, we measured ratios of transfer entropies in forward, backward and sideways directions for PDMS+frame (Fig 8 G), PDMS-frame (Fig 8 H) and for standard cultures (Fig 8 I). Unidirectionality is clearly seen in PDMS+frame cultures, but is less prominent in PDMS-frame cultures, where there’s mostly evidence for a bi-directional propagation of electrical activity. In standard cultures there is, as expected, no preferred direction for neuronal activity.

## Discussion

In this article, we’ve reported on a microtunnel design for producing artificial neuronal networks with population-wide control of neurite growth. We have shown that different patterns produce different growth morphology, and that these growth patterns can be reproduced in a simple simulation. By interfacing one of our microtunnel designs with microelectrode arrays, we could confirm that the morphological influence also extends to the electrophysiological network activity of the neurons. We also noted a serendipitous but highly significant increase in signal to noise ratio when using PDMS microtunnels on microelectrode arrays.

Using microfabricated tunnels to guide neurite growth has been used extensively in the past[37, 25, 10], but has been mostly confined to asymmetric connections between a small number of neuronal populations which grow outside the microtunnel compartment. While these experiments can be highly informative, for example by allowing the isolation of two or more cell types influence on each other [22, 16], controlling neuronal networks on a global scale opens up entirely new doors for what can be done in vitro. As mentioned, there’s been tremendous advancements in our ability to model health and disease of the human nervous system in vitro. Today, the use of embryonic stem cells and induced pluripotent stem cells have become widely adopted. Combined with 3D cultures, and the use of reprogramming and refined induction protocols[53], in vitro research continues to push the boundary of what is possible to recreate in vitro. Despite these impressive advances, almost all in vitro systems to date show highly synchronous electrical activity which isn’t normally found in a mature healthy brain, potentially limiting what phenomena can be effectively modeled. Controlling neural network architecture could be a way to circumvent this limitation and create neural network which better replicate in vivo networks.

A useful concept for describing neuronal networks in this context is the notion of integration and segregation. In order for a brain to separate different pathways, they must be segregated, while some degree of integration is important for the brain to make associations and exchange information between pathways[43, 37]. A typical in vitro neuronal culture is in this sense disproportionally integrative with very little segregation. This is evident by the fact that electrical stimulation in neuronal cultures tends to evoke activity across the entire culture[23, 17, 39, 36], while in vivo or ex vivo preparations show more confined responses to stimuli[7, 6, 5]. Our perforated microtunnel design could allow replication of the brains integrative and segregational structure.

Another important note here is that, while we are aware of a few experiments which utilize a similar approach to yield large quantities of neurons inside microtunnels, these tend to be either fully ‘integrating’ by allowing bidirectional communication[44], or entirely ‘segregational’ by strictly allowing only unidirectional connections to form[9, 30]. As mentioned, a brain express a combination of both integration and segregation[45], and our tunnel design offers the capacity to weight these two independently. One could conceivably use combinations of the microtunnel designs described here, with solid walls which completely segregates different parts of the network, and investigate if these can reveal nuances in neuronal activity which may otherwise go unnoticed in traditional ‘bursting’ cultures.

Using this technique to construct and investigate various network architectures in vitro is highly permissible with the approach we describe here, however, we found that neurons tend to grow in paths which were difficult to predict. Using our computer simulation could be a useful tool for designing complex network architectures, allowing a form of computeraided design (CAD) for in vitro neuronal networks.

An important limitation to this technique is that cells within perforations are not guided by the microtunnels, rather they likely form a highly interconnected network within each perforation. However, this is perhaps not too dissimilar to cortical columns of clustered network topologies which are common within the cortex[32]. Another limitation is that the microtunnels are not explicitly selective for axons, and dendrites may grow and form connections in the “wrong” direction. In similar settings, it’s usually assumed that dendrites do not grow as far as axons, thus enabling control of direction selectivity in neuronal communication[14].

## Conclusions

In conclusion, we have shown that our microtunnel design can serve as a powerful platform for guiding neuronal circuit formation in vitro. The design was readily interfaced to microelectrode arrays and application of cells was possible with no modifications to standard seeding protocol. The ability to guide the propagation of neuronal activity, as well as the reliable and great signal to noise characteristics, means that this design could serve as a useful tool when studying electrophysiological network properties of neuronal circuits in vitro.

## Acknowledgments

This work was funded by Vetenskapsrådet (2022-04760), Crafoordska Stiftelsen (20220951) and Carl Tryggers Stiftelse (CTS 23: 2839).

