## Supplementary Figure 1 for "Large-Scale Control of Neuronal Networks In Vitro Using Perforated Microfluidic Devices"

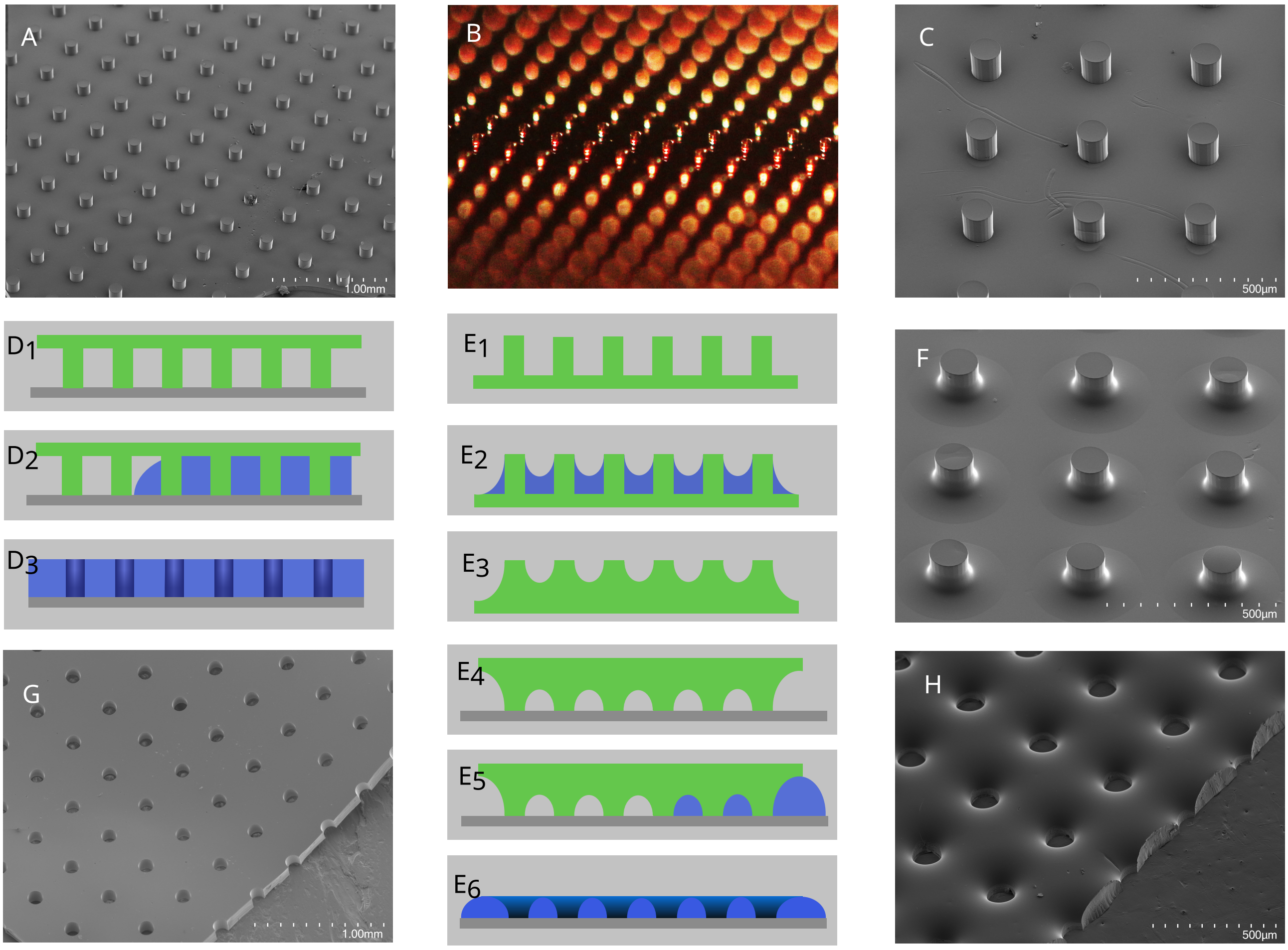


Figure 1: Production of different pillar and perforation morphology. A) Pillars of PDMS. B) Original wafer with pillars. C) Normal PDMS pillars. $D_1$) PDMS pillars are interfaced to a glass surface, which can be attached firmly by plasma-treating or heat-treating in 80 degrees. $D_2$) PDMS is flooded into the void between the pillars. $D_3$) Upon removing the PDMS pillars, a perforated membrane is created. $E_1$) Original PDMS pillars. $E_2$) A small volume of PDMS is added to smooth the pillars. $E_3$) PDMS cures and is left to bind to the pillars. $E_4$) The 'smoothed' pillars are interfaced to a glass substrate as in 'D'. $E_1$) PDMS is flooded into the void between the smoothed pillars. $E_1$) A 'dimpled' perforated membrane is created. F) Smoothed pillars. G) 'Flat' perforated membrane. H) 'Dimpled' perforated membrane.
